# IL-13 promotes recovery from *C. difficile* infection

**DOI:** 10.1101/2021.08.03.454972

**Authors:** A.N. Donlan, J.L. Leslie, M.E. Simpson, W.A. Petri, J.E. Allen, W.A. Petri

## Abstract

*Clostridioides difficile* infection (CDI) is the leading hospital-acquired infection in North America. We have previously discovered that antibiotic disruption of the gut microbiota decreases intestinal IL-33 and IL-25 and increases susceptibility to CDI. We further found that IL-33 promotes protection through type 2 Innate Lymphoid Cells (ILC2s), which produce IL-13. However, the contribution of IL-13 to disease has never been explored. We found that administration of IL-13 protected, and anti-IL-13 exacerbated CDI as measured by weight loss and clinical score, particularly during disease resolution. Additionally, concordant with IL-13 being important for M2 macrophage polarization, we saw a decrease in M2 macrophages (CD11B+CD64+CD206+) cells following neutralization of IL-13. We also observed monocyte accumulation as early as day three post-infection following IL-13 neutralization, suggesting IL-13 may be directly or indirectly important for their recruitment or transition into macrophages. Neutralization of the decoy receptor IL-13Rα2 resulted in protection from disease, likely through increased available endogenous IL-13. Our data highlight the protective role of IL-13 in promoting recovery from CDI and the association of poor responses with a dysregulated monocyte-macrophage compartment. These results increase our understanding of type 2 immunity in CDI and may have implications for treating disease in patients.

## Introduction

*Clostridioides difficile* infection (CDI) is the leading hospital-acquired infection in North America. Infection with *C. difficile* is most often associated with antibiotic usage resulting in the disruption of normal host microbiota. The host immune response to infection is an important driver of disease outcomes, as white blood cell (WBC) count above 20 x 10^9^/L and high levels of inflammatory markers are better indicators of severe disease than bacterial burden^1,2^. While antibiotics are available to treat patients with CDI, there were 29,000 deaths reported in 2011, and 1 in 5 will still progress to relapsing disease^3^, highlighting the need for alternative therapeutics.

Limiting pathogenic inflammation in hosts may be a novel approach to protecting against severe outcomes. We have previously shown that administration of recombinant IL-33 during murine CDI provides protection from severe disease, and that this was facilitated by type 2 innate lymphoid cells (ILC2s)^4^. During disease, ILC2s from IL-33-treated mice expressed increased levels of IL-13, implicating this cytokine in mediating protection from IL-33, but the role of IL-13 during CDI has not been explored.

IL-13 is an important effector cytokine for type 2 immune responses and has roles in driving polarization of alternatively-activated macrophages^5,6^, decreasing Th17-associated inflammation^7,8^, and promoting eosinophil recruitment as well as goblet cell hyperplasia^9^. As our lab has previously described, the closely related cytokine to IL-13, IL-4, was increased following administration of IL-25, and neutralization of IL-4 resulted in delayed, while administration promoted, recovery in mice^10^. Because of the similarity in functions to IL-4, we hypothesized that IL-13 would be important for during resolution from infection with *C. difficile*. Through neutralization and administration of IL-13, we observed that this cytokine was associated with promoting recovery from disease. We also observed that IL-13 neutralization was associated with increased Ly6C-high monocytes and reduced CD206+ macrophages, suggesting that these cells may impede or promote recovery responses, respectively. Together, our data shed light on another important contribution of type 2 immunity to protection from infection with *C. difficile*.

## Results

To test the role of endogenous IL-13 during CDI, we infected mice with the R20291 spores of *C. difficile* and administered i.p. injections of anti-IL-13 neutralizing antibodies or IgG control on days 0 and 2 post infection. Mice who received anti-IL-13 antibodies developed disease similar to control, but had delayed recovery by clinical scores and weight (**Figure 1A**). In contrast, administration of recombinant IL-13 promoted recovery in mice (**Figure 1B**). Neither neutralization or administration had significant impact on mortality due to disease (**Figure 1, C and D**). Together, however, this implicated the importance of IL-13 in promoting recovery in a murine model of CDI. Additionally, *C. difficile* burden was not different between IL-13 neutralized mice and controls on either day three or four post infection (**Figure 1E**), suggesting that changes to bacterial burden was not driving differences in recovery outcomes.

**Figure 1.**
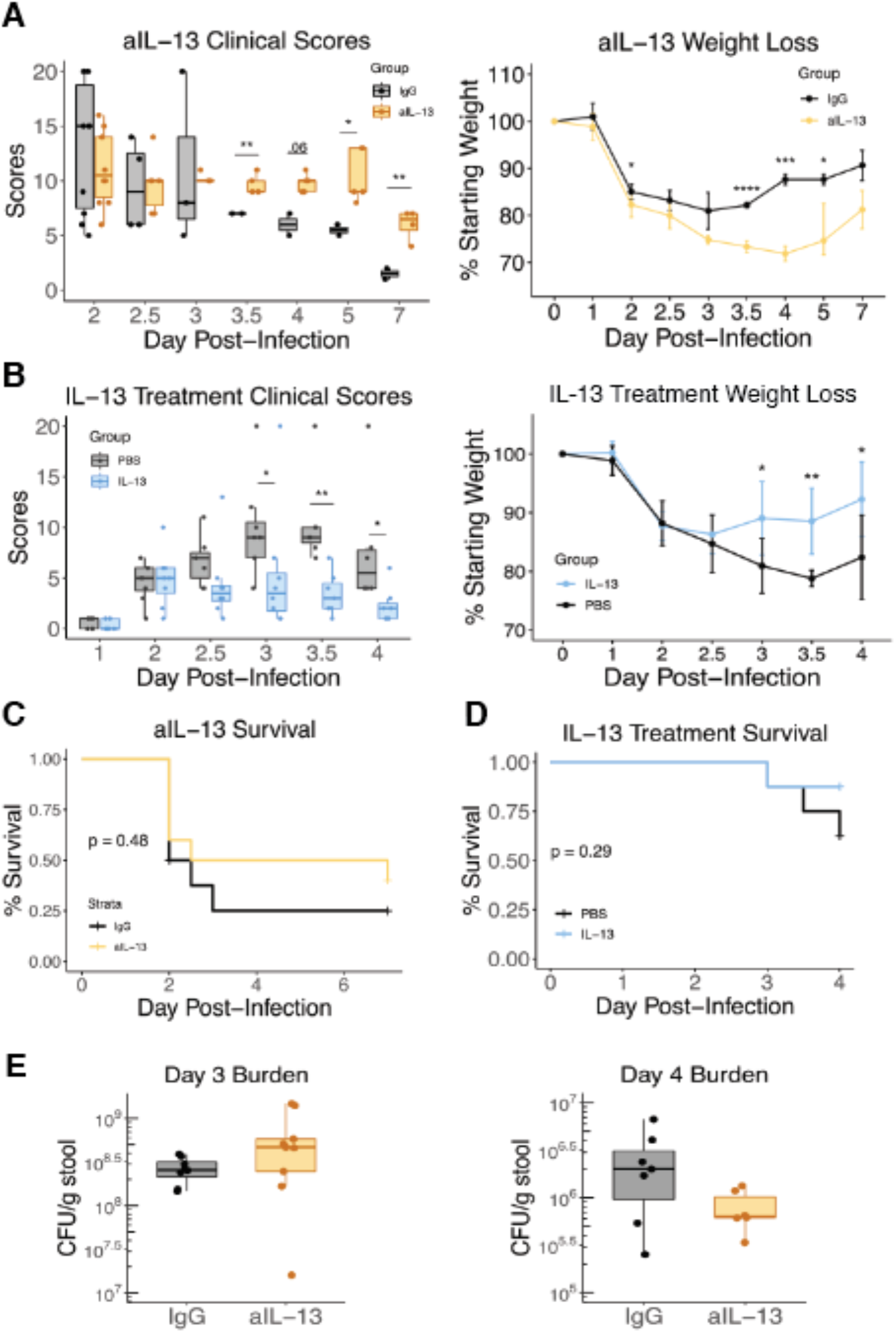
IL-13 provides protection from *C. difficile* infection. A) Weight loss and clinical scores for mice i.p. injected with 150 μg of anti-IL-13 antibodies or IgG control on days 0 and 2 post-infection with 100-1000 spores of *C. difficile*. B) Weight loss and clinical scores of mice i.p. injected with 100 μL of 5 μg of recombinant IL-13 or PBS control. C,D) Kaplan-Meier curves for treatments in A) and B). E) *C. difficile* burden on days 3 and 4 post-infection. * = p<0.05; ** = p<0.01; *** = p <0.001. Student t-test.

Because overabundant immune responses during CDI have been shown to be deleterious for host outcomes, we tested whether the neutralization of IL-13 was impeding the resolution of inflammation, resulting in delayed recovery. By day four postinfection, when mice typically would begin recovering, neutralization of IL-13 resulted in increased inflammation within the lamina propria. There were elevated levels of total immune cells (**Figure 2A**), as well as inflammatory myeloid cells such as neutrophils and monocytes (**Figure 2, B and C**). Because IL-13 can have an important role in promoting the polarization of alternatively-activated, CD206+, macrophages (AAMs)^6,11^, which can be important for anti-inflammatory processes and tissue repair, we wanted to test whether changes to these cells due to neutralization preceded the increased inflammation seen on day 4. On day three post-infection, there were no differences in total number or proportion of CD64+ (FcGR1) macrophages^12^ in the lamina propria, but there were significantly reduced CD206+ macrophages (**Figure 2E)**. Notably, when gating on CD64+ cells we observed that there was a significant increase in a population of cells expressing low levels of this marker (**Figure 2F**). Using a waterfall plot, which shows the transition from Ly6C high monocyte to MHCII+ tissue macrophage^12^, we observed that these CD64-low cells were mainly LyC6-high MHCII low/-, suggesting that these cells represent monocytes that are beginning to differentiate into tissue macrophages. In contrast, CD64-high cells appeared to be comprised of cells further along that transition pathway from LY6C+ MHCII- monocytes to Ly6C-MHCII+ macrophages. Accumulation of these CD64-low, transitionary, cells then, could be impeding recovery by contributing to increased inflammation. Additionally, this could suggest the reduced capacity of monocytes to transition into tissue macrophages due to potentially increased inflammatory environment. In support of this, although there was no change in CD64+ cells on day three post-infection, we observed a trend towards decreased CD64+MHCII+ macrophages (**Figure S1)** by day 4 post-infection. Together, these data support the hypothesis that there are dysregulated monocyte and macrophage responses following IL-13 neutralization.

**Figure 2.**
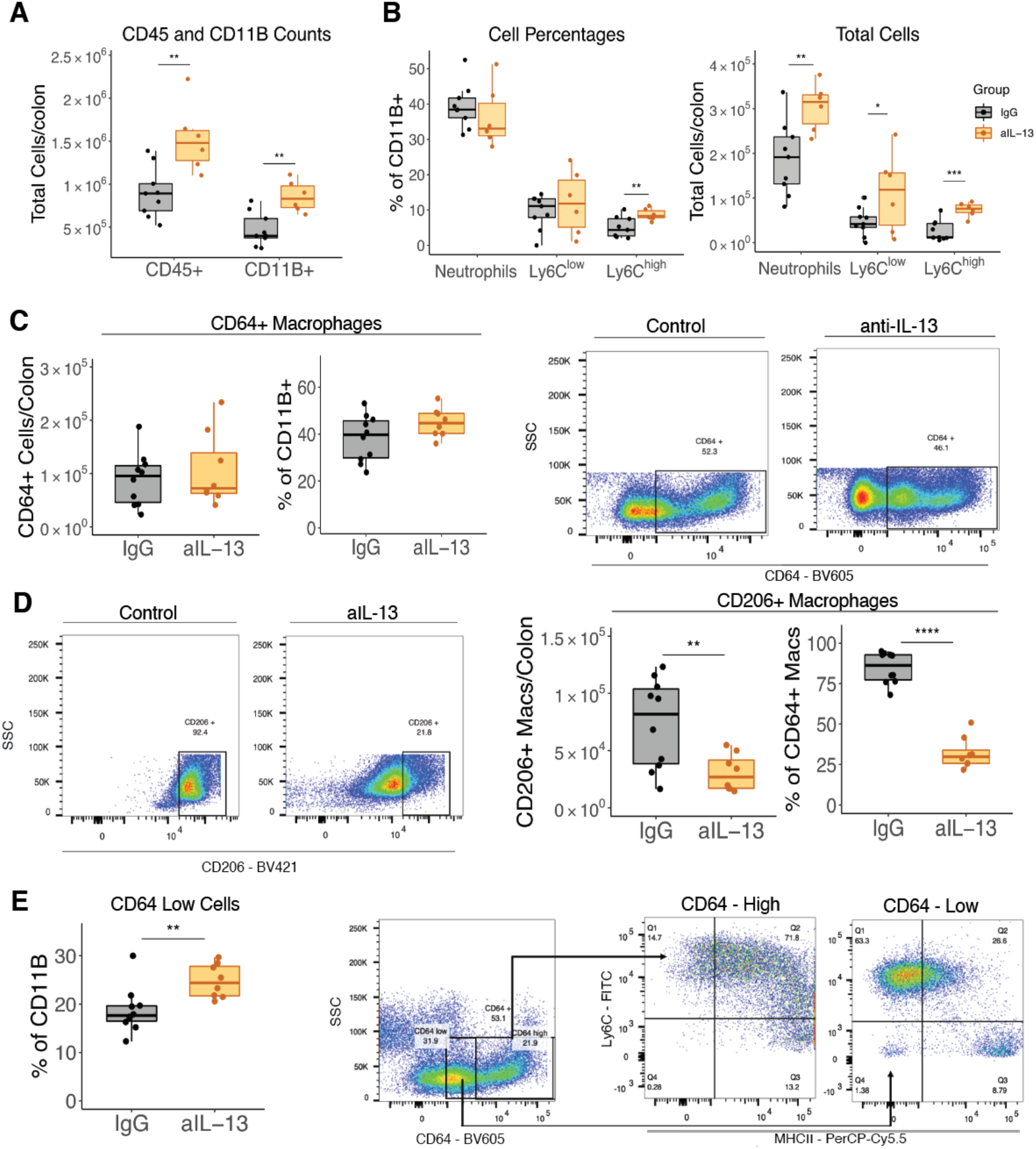
Neutralization of IL-13 increases inflammation and monocyte accumulation while decreasing abundance of CD206+ macrophages in the colon. Mice were treated anti-IL-13 (orange) or IgG (grey) antibodies on days 0 and 2 postinfection with *C. difficile*. A) Total number per colon of CD45+ and CD11B+ cells on day four post-infection. B) Total number per colon and percent of CD11B+ myeloid cells for CD11B+Ly6G+Ly6Clow neutrophils, CD11B+Ly6G-Ly6C -high and -low monocytes on day four post-infection. C) Percent of CD11B+ cell and total cells per colon of CD11B+CD64+ macrophages on day three post-infection. D) Representative flow plots of (C). E) Percent of CD64+ macrophages and total cells per colon of CD11B+CD64+CD206+ macrophages on day three post-infection. Representative images below. F) Percent of CD11B+ cells of CD11B+CD64-low cells on day 3 postinfection. G) Representative flow plot and waterfall plot (Ly6C vs MHCII) of CD64-low and CD64-high gated cells.. * = p<0.05; ** = p<0.01; *** = p <0.001. Student t-test.

IL-13 can promote the expression of IL-10 in AAMs^13^, and accordingly, treatment with IL-13 also resulted in increased IL-10 expression by day 5 post-infection (**Figure S2**). IL-10 is a potent anti-inflammatory cytokine, and this suggests that IL-10 may be important for IL-13-mediated recovery responses. Because CD206+ AAMs have been implicated in the recovery and resolution of other inflammatory insults, we wanted to observe whether they corresponded to the recovery process of CDI. Using flow cytometry, we observed the myeloid compartment in the lamina propria of mice on days two, four, and five post infection, compared to uninfected mice with or without antibiotics (**Figure 3, A and B**). As expected, we saw an increase in inflammatory cells such as neutrophils and monocytes following infection. CD206+ macrophages were predominant in the uninfected colon, but expanded further by day 5 post-infection, suggesting they are associated with the recovery phase of disease (**Figure 3C**).

**Figure 3.**
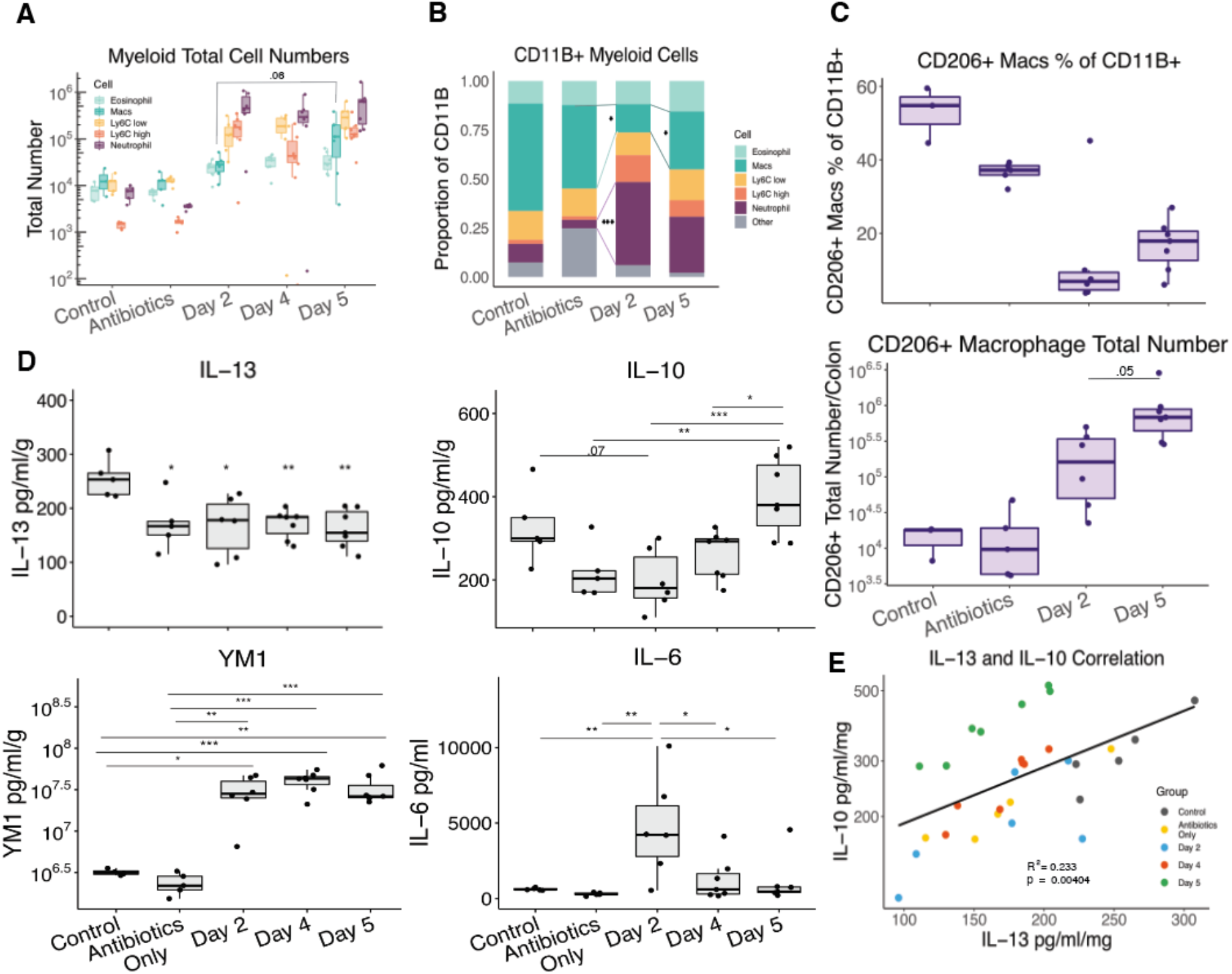
Recovery from *C. difficile* is associated with macrophages and antiinflammatory cytokines. Mice were left as uninfected controls, or given normal antibiotic regimen with or without subsequent infection with 500 spores of R20291 *C. difficile*. Mice were euthanized on day 2, 4 and 5 post-infection. A, B) Total number per colon and percent of CD11B+ cells of neutrophils, monocytes, eosinophils, and CD64+ macrophages. C) Total number and percent CD11B+ cells of CD64+CD206+ macrophages. D) Cecal tissue was lysed and IL-13, IL-10, YM1, and IL-6 ELISAs were performed. E) Scatter plot showing correlation between IL-13 and IL-10. Colors indicate which group samples belong to. Linear regression and p-value shown. Tukey’s HSD; * = p<0.05; ** = p<0.01; *** = p <0.001. Student t-test.

Next, we measured IL-13 levels during the disease course, but did not observe any increase in expression by ELISA (**Figure 3D**). The lack of increased protein, however, could still be resulting in increased signaling through potentially elevated receptor expression. IL-10, in contrast, was increased during disease recovery and was significantly correlated with IL-13 levels (**Figure 3, D and E**). YM1, a type 2 marker, was also increased following infection, supporting data from Frisbee et al. Additionally, as an indicator that a pro-inflammatory mediator was associated with acute disease as opposed to recovery, we measured IL-6, which was most highly abundant on day two post-infection (**Figure 3D**).

Because CD206+ macrophages are typically monocyte-derived^14^, to test whether CCR2+ monocyte-derived macrophages are important for infection, we depleted these cells using anti-CCR2 antibodies^15^ on day two post infection. Loss of monocytes at this time point resulted in worsened disease, suggesting an important contribution during host immunity (**Figure 4A**). However, this approach would additionally deplete monocytic lineages not dependent on IL-13, which may account for the large impact on survival outcomes. In contrast, monocyte depletion at earlier time points resulted in no change to disease (**Figure 4B**) which suggests that early monocytes give rise to inflammatory cells that are balanced by anti-inflammatory cells generated at subsequent time points.

**Figure 4.**
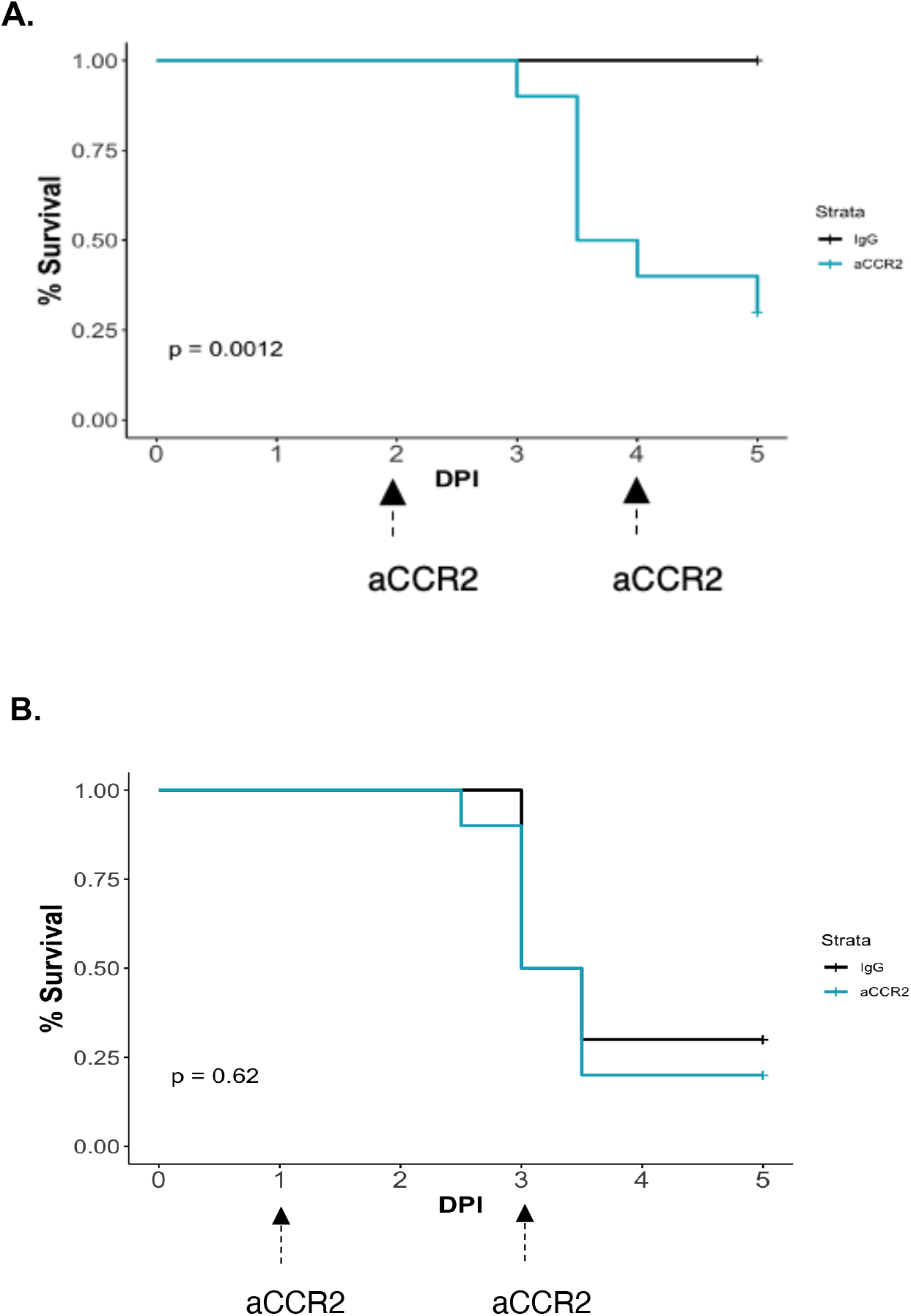
Depletion of monocytes at later timepoints worsens disease. Mice were administered 500 μg of anti-CCR2 depleting antibodies on days A) 2 and 4, or B) 1 and 3 post-infection. Kaplan-Meier curves shown (log rank test).

The decoy receptor for IL-13, IL-13Rα2, has been shown to be important for recovery in a DSS colitis animal model^16^, and as such, we wanted to test whether this negative regulator of IL-13 could be involved in CDI as well. We measured levels of this receptor during infection by ELISA and observed that it increased during the recovery phase of disease (**Figure 5A**). Since the expression of IL-13Rα2 is positively regulated by increased IL-13 signaling as a method for negative feedback, this supports the hypothesis that IL-13 could be contributing during the recovery process. Additionally, IL-13Rα2 can be positively regulated by other proinflammatory factors such as TNFα and IL-17^17^. Since both of these cytokines are increased during CDI in the animal model, this may indicate that the inflammatory response initiated by *C. difficile* infection is poised to dampen a type 2 immune response mediated by IL-13. IL-13Rα2 was also negatively correlated with weight (**Figure 5B**), further suggesting that high levels are not beneficial to the host. To test the contribution of IL-13Rα2 to CDI, we administered neutralizing antibodies on days 1 and 3 post-infection. IL-13Rα2 neutralization resulted in improved weight and clinical scores, indicating that this decoy receptor negatively regulates protective type 2 immunity through IL-13 (**Figure 5C and D**).

**Figure 5.**
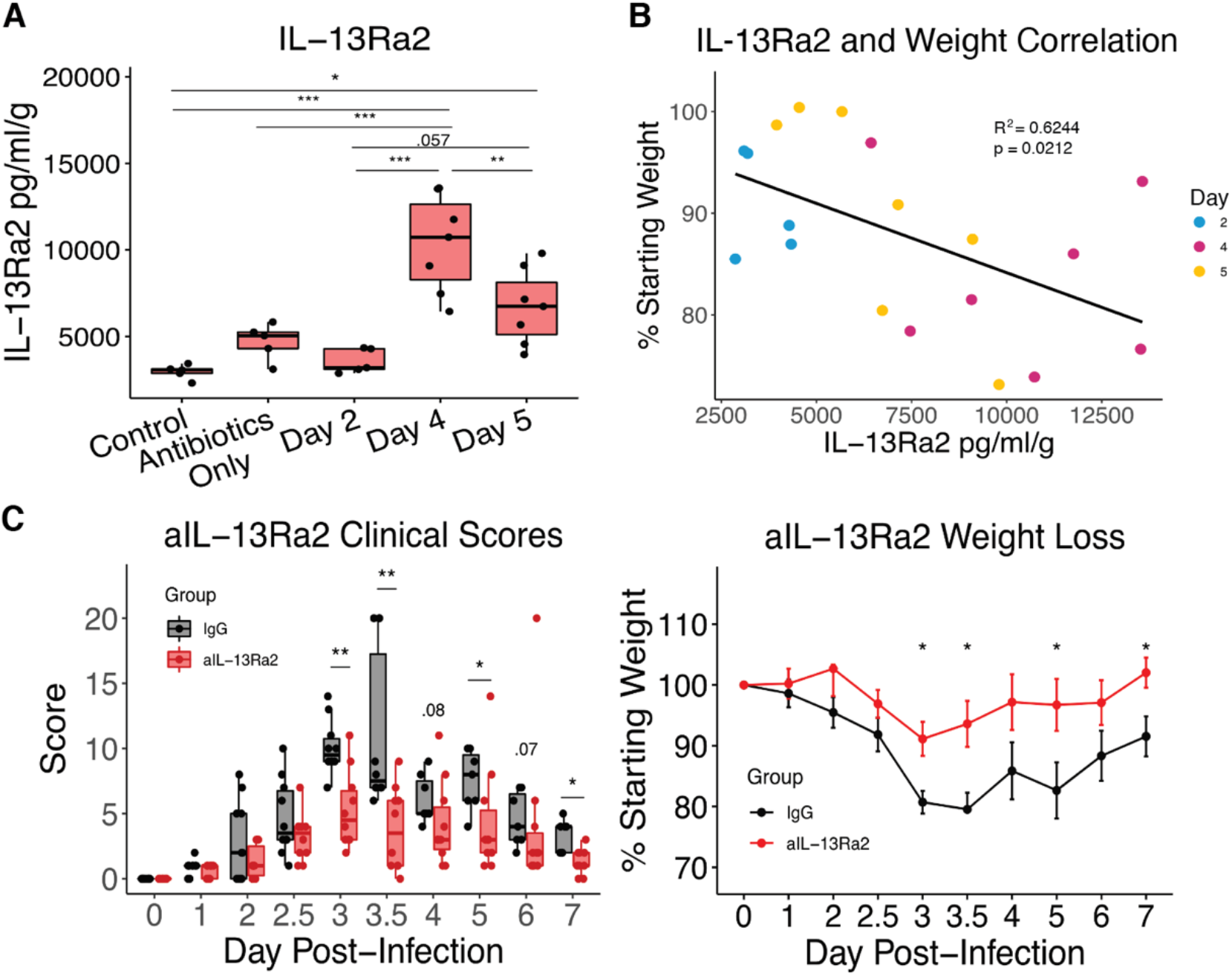
IL-13Rα2 neutralization protects during CDI. A) IL-13Rα2 protein levels in cecal lysates from from uninfected mice with or without antibiotic administration, or mice on days 2, 4, and 5 post infection were tested by ELISA and normalized to tissue weight (Tukey’s HSD). B) Linear regression of IL-13Rα2 levels with weight loss (Spearman). C,D) Mice were injected with 250 μg of anti-IL-13Rα2 or IgG1 control on days 2 and 4 post infection. C) Clinical scores and D) weight loss (T-test).* = p < 0.05; ** = p < 0.01; *** = p < 0.005.

## Discussion

The importance of type 2 immune responses in protecting against pathogenic type 17 inflammation following *C. difficile* infection is further supported here, by the evidence that IL-13 contributes to recovery from infection.

IL-33 was previously shown to be protective during CDI through the induction of ILC2s, which in turn produced higher levels of IL-13 following IL-33 treatment. However, the contribution of IL-13 during CDI had not yet been explored. Here we report that neutralization or administration of IL-13 during a murine model of CDI had an impact on the recovery phase of disease, suggesting that IL-13 contributes during this response.

IL-13 can be an important regulator of inflammation and is often considered an anti-inflammatory cytokine outside the context of pathogenic type 2 inflammation such as asthma. Importantly, IL-13 contributes to the polarization of CD206+ alternatively-activated macrophages, which are generally associated with anti-inflammatory functions and tissue repair^7,9,11^.

In our model, when IL-13 was neutralized, we observed a decrease in CD206+ macrophages by day four post infection, implicating them as potential downstream mediators of IL-13 functions. We observed the highest levels of these CD206+ macrophages during the recovery phase of disease, highlighting that they may be involved in this process. Furthermore, we observed that neutralization of IL-13 resulted in increased Ly6C-high monocytes that accumulated in a transitionary phase prior to becoming tissue macrophages. Whether this is due to increased chemokine expression, or if increased inflammation reduces the capacity for monocytes to transition into macrophages, is unknown. IL-13 has been observed to have roles in reducing CCL2 /monocyte-chemoattracting protein −1 (MCP-1) expression in some contexts^18,19^, resulting in decreased monocyte recruitment from the bone marrow. However, increased inflammation in the colon during DSS colitis was also associated with an accumulation of Ly6C-high monocytes^20^, suggesting that inflammation may have an impact on monocyte-derived macrophage development or accumulation of monocytes within the tissue.

Although IL-13 was induced downstream of IL-33, which protected mice from mortality and severity during acute disease, our data suggested that IL-13 was not the primary facilitator of the acute protection conferred by IL-33. However, it is likely that IL-13 mediates many of the responses important for reducing inflammatory cells within the tissue. The potential for this is highlighted by the increase in immune cells by day four post infection following IL-13 neutralization, suggesting the environment is more inflammatory.

We have previously shown that IL-4 is involved in the recovery from *C. difficile* infection^10^. Given that IL-4 and IL-13 share a high degree of similarity in sequence, signaling, and influence on immune responses, it would be reasonable to hypothesize that these cytokines have overlapping functions during CDI. If one can compensate for the other, when absent, it is possible that dual neutralization or administration may uncover a larger contribution for this signaling pathway during infection.

Overall, this work increases our understanding of the protective capabilities for type 2 immunity during *C. difficile* infection, and may have broader implications for other enteric inflammatory responses.

## Acknowledgements

This work was supported by the National Institutes of Health (grant numbers R01 AI124214 to WAP). Anti-IL-13Ra2 was a gift from Dr. Thomas A. Wynn. Thomas Vasquez performed IL-10, IL-6, and CCL2 ELISAs.

## Author contributions

A.N.D. conceived, designed, and performed all experiments and wrote the manuscript. J.L.L and M.E.S assisted in tissue processing and invaluable discussion. W.A.P. assisted with technical support and running of many experiments. J.E.A. provided guidance and advice. W.A.P. Jr supported all aspects of the work

## Methods

### Bacterial strains and culture

To generate spore stocks, vegetative *C. difficile* strain R20291^21^ [3] was plated on BHI agar from frozen stocks and incubated overnight at 37°C in an anaerobic chamber (Shel Labs). A single colony was then inoculated into 15 mL of Columbia broth overnight at 37°C anaerobically. Then 5 mL of turbid broth was moved into 45 mL of Clospore broth^22^ and incubated at 37°C for 5-7 days. Cultures were spun down at 3200 rpm for 20 minutes at 4°C then resuspended in cold sterile water. This was repeated 3-5 times to lyse vegetative cells. The pellet was then resuspended in 1 mL sterile water and transferred to a 1.5 mL twist cap tube (Corning # 4309309). Spores were plated on BHIS +Taurocholate agar plates to determine concentration of stock. Spores were stored at 4°C. To make the inoculum, a 5×10^4^-1×10^5^ spores/mL dose was prepared in sterile water. Each mouse received 100 μL (5×10^3^ −5×10^4^ spores) by oral gavage. *C. difficile* burden was quantified from cecal contents following infection. Total cecal content was resuspended by weight in sterile, anaerobic PBS. Serial dilutions of cecal contents were plated on BHI agar supplemented with 1% taurocholate, 1 mg/mL D-cycloserine, and 0.032 mg/mL cefoxitin (Sigma) and incubated anaerobically at 37°C overnight before colonies were counted.

### Mice

Experiments were carried out using male, 8-12-week-old C57BL/6J mice ordered from Jackson Laboratory. All animals were housed under specific pathogen-free conditions at the University of Virginia’s animal facility. Mice were infected following a previously published murine model of CDI^4,21^. Six days prior to infection mice were given a cocktail of antibiotics in their drinking water, consisting of 45 mL/L Vancomycin (Mylan), 35 mg/L Colistin (Sigma), 35 mg/mL Gentamicin (Sigma), and 215 mg/L Metronidazole (Hospira). Mice were then switched to normal drinking water for three days prior to infection. One day before infection, mice were given an IP injection (0.016mg/g) of Clindamycin (Hospira). On the day of infection, mice were orally gavaged with 5×10^3^ −5×10^4^ spores of *C. difficile* strain R20291 in 100 μL water. Mice were monitored twice daily over the course of infection and evaluated according to clinical scoring parameters. Scores were based on weight loss, coat condition, activity level, diarrhea, posture, and eye condition for a cumulative clinical score between 1 and 20; scores were not blinded. Weight loss and activity levels were scored between 0 and 4, with 4 being greater than or equal to 25% loss in weight. Coat condition, diarrhea, posture, and eye condition were scored between 0 and 3. Diarrhea scores were 1 for soft or yellow stool, 2 for wet tail, and 3 for liquid or no stool. Mice were euthanized if severe illness developed based on a clinical score ≥ 14. On the day of and one day post infection, For IL-13 treatment, mice were i.p. injected with 5 μg of recombinant murine IL-13 (Peprotech cat# 210-13) in PBS (100 μL dose) on days −1 and 1 of infection. Control treatments for all groups were filter-sterilized PBS. For anti-IL-13 neutralization, mice were administered 150 μg of anti-IL-13 (eBio1316H cat# 16-7135-85) or IgG (eBRG1 cat# 16430185) from Thermofisher. For anti-CCR2 cell depleting treatment, mice were administered 500 μg of anti-CCR2 (MC-21; gifted from Dr. Matthias Mack) on either days 2 and 4, or 1 and 3. Anti-IL-13Rα2 treatment was administered using 250 μg of anti-IL-13Rα2 (clone: 6D5) or IgG1 (HPRN) (gifted from Dr. Tom Wynn) on days 2 and 4 post-infection.

### Flow cytometry

Colons were dissected longitudinally and rinsed in HBSS (Thermofisher) supplemented with 25 mM HEPES and 5% FBS. Epithelial cells were separated from the lamina propria via a 40-min incubation with gentle agitation in dissociation buffer (HBSS with 15 mM HEPES (Thermofisher, 5 mM EDTA (Invitrogen), 10% FBS (Gibco), and 1 mM DTT (Bioworld)) at 37 °C. Next, the lamina propria tissue was manually diced using scissors and further digested in RPMI 1640 (Gibco) containing 0.17 mg/mL Liberase TL (Roche) and 30 μg/mL DNase (Sigma). Samples were digested for 40 min at 37 °C with gentle shaking. Single-cell suspensions were generated by passaging samples through a 100 μM cell strainer followed by a 40 μM cell strainer (Fisher Scientific). For staining, single-cell suspensions were obtained and cells were stained with the following monoclonal antibodies: CD45 (APC-Cy7 #103116 1/200), CD11B (APC #101212, 1/200), Ly6G (PE-Cy7 #127618, 1/100) from Biolegend, Siglec-F (PE #552126, 1/100) from BD Biosciences, CD64 (BV711 clone: X54-5/7.1 #139311), CD206 (BV421 # 141717) from Biolegend, MHCII (PerCP-Cy7 clone: M5/114.15.2 #107626) from Biolegend, F4/80 (FITC clone: BM8 #123108) from Biolegend, IL-10 (PE-Dazzle #505033) from Biolegend, Arg1(PE # 12-3697-82) from Thermofisher. For surface staining, 1 × 10^6^ cells/sample were Fc-blocked with TruStain fcX (BioLegend, #101320, 1/200) for 10 minutes at room temperature followed by LIVE/DEAD Fixable Aqua (Life Technologies) for 30 min at 4 °C. Cells were washed twice in FACS buffer (PBS+ 2% FBS) and stained with fluorochrome conjugated antibodies for 30 min at 4 °C. Cells were washed twice before incubation for 20 minutes at 4 °C in BD Cytofix (BD Cytofix/Cytoperm Fixation/Permeabilization Solution Kit, #555028) and then were washed and resuspended in FACS buffer and incubated overnight at 4 °C. Flow cytometry was performed on an LSR Fortessa cytometer (BD Biosciences) and all data analysis performed via FlowJo.

### ELISAs

R7D Mouse Duoset Sandwich ELISA kits were used to measure levels of IL-13 (cat# DY413-05), IL-10 (cat# DY417-05), YM1 (CHI3L3) (cat# DY2446), IL-6 (cat# DY406-05), IL-13Rα2 (cat# DY539), and Hyaluronan (cat# DY3614-05) were detected in cecal tissue lysates according to manufacturer’s instructions. Total cecal lysate was generated by removing the ceca and rinsing gently with 1x PBS. Tissue was bead beat for 1 minute and resuspended in 400 μL of Lysis Buffer I: 1× HALT Protease Inhibitor (Pierce), 5 mM HEPES. Following mechanical tissue disruption, 400 μL of Lysis Buffer II was added: 1× HALT Protease Inhibitor (Pierce), 5 mM HEPES, 2% Triton X-100. Tissue samples were incubated on ice for 30 minutes after gently mixing. Lysed samples were pelleted to remove tissue debris in a 5-minute spin at 13,000 × *g* at 4°C. Supernatant was collected and total protein concentration was measured by BCA assay according to manufacturer’s instructions (Pierce). Cytokine concentration was normalized to tissue weight.

### Statistical Analyses

Survival differences between groups were tested using the log-rank test. Statistical differences in clinical scores and weight loss were evaluated using Student’s t-test (twotailed). Linear regressions were performed in R studio. A p-value below 0.05 was considered significant. All statistical tests were run using R Studio software (Version 1.2.1335).

## Supplement

**Figure S1.**
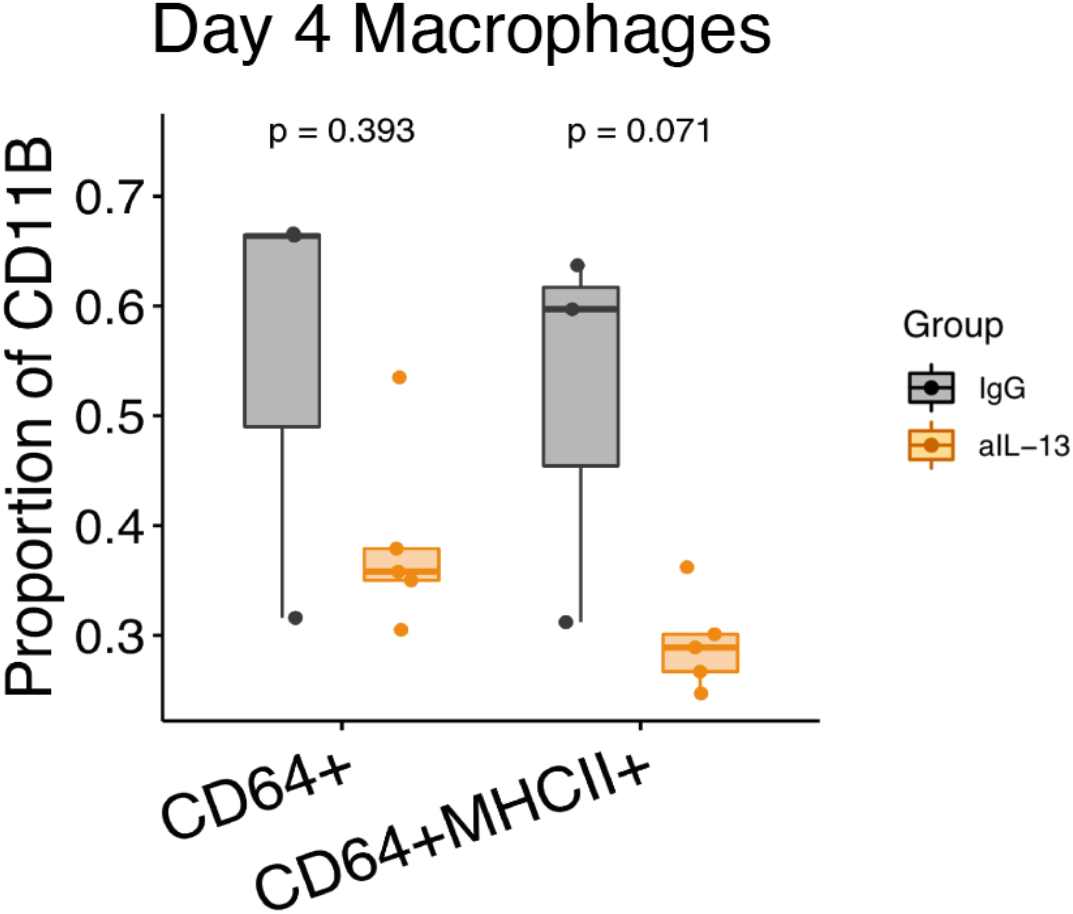
IL-13 neutralization results in trend towards decreased macrophages by day 4 post infection. Mice were treated anti-IL-13 antibodies on days 0 and 2 following infection with *C. difficile* and colonic tissue was taken on day four postinfection. Percent of CD11B for CD64+ and CD64+MHCII+ cells between IgG and anti-IL-13 treated mice. T-test; p values shown. N = 7.

**Figure S2.**
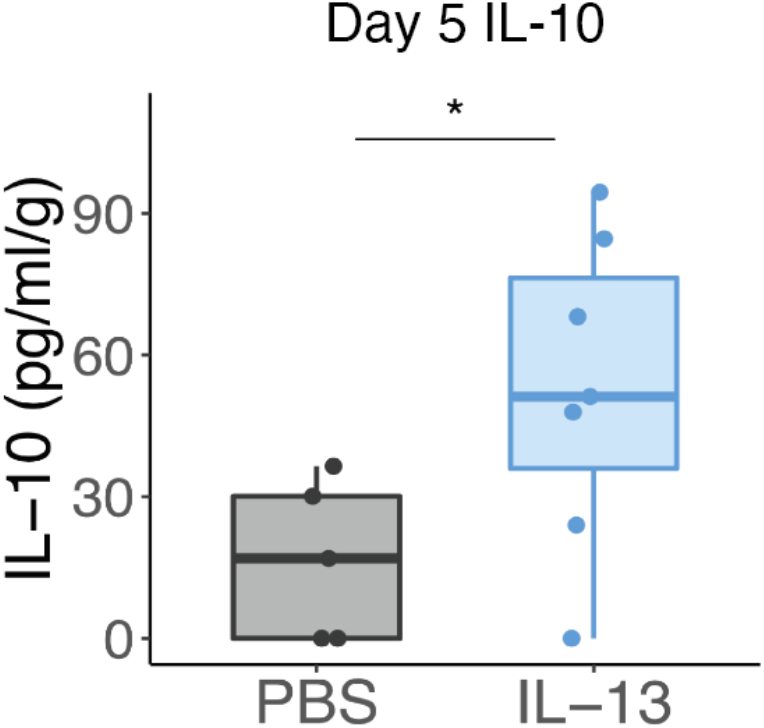
IL-13 treatment alters IL-10 on day 5 post-infection. Mice were i.p. injected with 100 μL of 5 μg recombinant IL-13 or PBS control on days 0, 1, and 2 postinfection with *C. difficile*. Cecal tissue was taken on day 5 post-infection and IL-10 levels were measured by ELISA and normalized to tissue weight. T-test; * = p < 0.05. N = 12.

